# Multi-dimensional DNA nanostructures isothermally assembled in hydrated ionic liquids

**DOI:** 10.64898/2026.09.02.744742

**Authors:** Hannah Talbot, Lauren Nguyen, Dhanush Gandavadi, Zohreh Nowzari, Chetna Mathur, Gwenyth Gallagher, Katrine Mongin, Sweta Vangaveti, Thomas J. Begley, Xing Wang, Arun Richard Chandrasekaran

## Abstract

DNA nanostructures can be tailored to perform a wide variety of functions, with continued interest in biological applications. Some aspects of DNA nanostructure assembly can hinder the ability of nanostructures to be useful in physiological environments. Typical assembly methods employ magnesium ions to stabilize the structure, which leave the structure susceptible to damage by nucleases in body fluids. Further, DNA nanostructure assembly typically involves a thermal annealing protocol in which DNA strands are heated in a specific buffer to a high temperature and cooled slowly at specific rates, preventing convenient encapsulation of temperature-sensitive guest molecules. In this work, we demonstrate the assembly of a wide variety of DNA nanostructures and 3D crystals in a hydrated ionic liquid (choline dihydrogen phosphate, CDHP) instead of magnesium at constant moderate temperatures, thus avoiding thermal annealing. CDHP-assembled structures show enhanced biostability against a variety of nucleases. Molecular dynamics simulations show that choline ions stabilize DNA nanostructures by a direct and close-range interaction in contrast to the predominantly water-mediated interactions of Mg^2+^, leading to enhanced nuclease resistance in CDHP-containing environments. CDHP-assembled structures do not affect the viability of HepG2 cells and show higher cell internalization. Overall, this work develops a potential method to construct more biostable DNA nanostructures and 3D crystals in a simple one-tube process. Assembly of DNA nanostructures under isothermal conditions is desirable for scaffolding biomolecules and to reduce the need for thermal annealing instruments, allowing nanostructure preparation in low-resource settings.

## Introduction

Through programmable design, DNA can be folded into finite shapes, periodic arrays and reconfigurable devices, with applications in biomolecular analysis, targeted delivery of drugs, data encryption, and diagnostics.^1–4^ For several applications, the desired robustness of these structures can vary, thus requiring different assembly conditions. While conceptual advances in DNA nanostructure self-assembly have allowed the construction of several complex structures and lattices,^5–8^ assembly is typically performed in buffers containing magnesium. DNA nanostructures can have limited stability in low-Mg^2+^ buffers, in nuclease rich environments and at elevated temperatures.^9–11^ To expand the assembly conditions as well to enhance the properties of DNA nanostructures, assembly has been studied in solutions containing other metallic counter ions,^12–14^ polyamines,^15,16^ cationic peptides,^17,18^ chaotropic agents,^19^ and organic solvents^20^ and compounds^21^ for ion-free self-assembly. In addition, the assembly of DNA nanostructures has been shown in atypical conditions such as deep eutectic solvents,^22^ but such methods require extreme conditions such as vacuum.^23^ Ionic liquids provide another alternative for such assemblies. For example, DNA and RNA aptamers have been assembled in ionic liquids such as choline dihydrogen phosphate (CDHP), which improved the biostability of these nucleic acids without impacting their function.^24^ In a recent work by our groups, we showed for the first time that DNA nanostructures can be assembled by thermal annealing in CDHP, further demonstrating higher biostability and enhanced functionality to target specific cell receptors.^25^

In typical DNA nanostructure self-assembly, the process involves a thermal annealing method, where the DNA solution is heated to high temperatures of 80-95 °C to eliminate secondary structures and gradually cooled down to a lower temperature over time. This annealing process is usually optimized to best support hybridization into the desired structure.^26^ However, the high temperatures used in the typical annealing process may not be compatible with every application, particularly ones utilizing guest molecules. To address these challenges, strategies for isothermal assembly of DNA nanostructures have been developed, where component DNA strands are mixed and incubated at a constant, more moderate temperature to construct nanostructures.^27^ The specific incubation temperature and the choice of counter ions can significantly affect the assembly of DNA nanostructures,^28–31^ thus necessitating the exploration of other ionic conditions for isothermal DNA self-assembly.^27^ Here, we demonstrate the isothermal assembly of DNA nanostructures in the hydrated ionic liquid CDHP. The choice of using hydrated versions of these ionic liquids allows ease-of-use with physiological applications,^32^ rapid folding of DNA nanostructures compared to water-free ionic liquids,^22^ and provides long-term stability for DNA.^33^ Further, the structural integrity of DNA nanostructures is based on the complexity of the structure, sequence design, and crossover density, among other parameters, necessitating the study of a variety of DNA nanostructures for a specific assembly condition.^34–36^ Our method allows the construction of DNA nanostructures (motifs and origami) as well as designed 3D crystals with enhanced properties provided by assembly in CDHP while allowing low resource assembly via isothermal method (**Figure 1**). Using MD simulations, we also provide mechanistic insights into the interaction of CDHP with a model DNA nanostructure.

**Figure 1.**
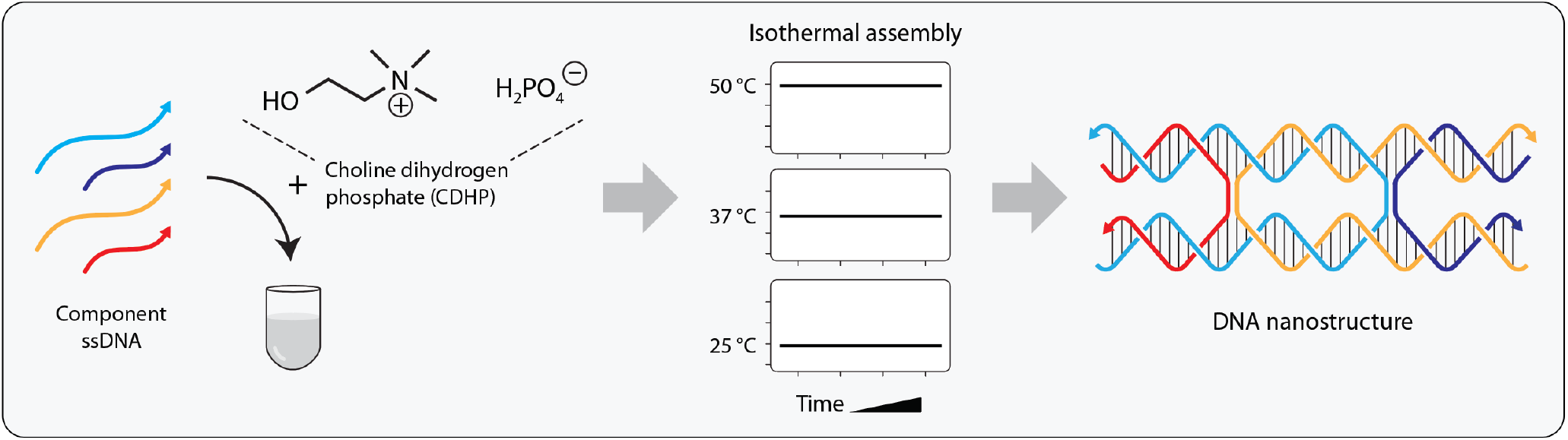
Isothermal assembly in CDHP. Component DNA strands are mixed in buffer containing choline dihydrogen phosphate (CDHP) instead of Mg^2+^ and incubated at a constant temperature of 25, 37 or 50 °C to assemble DNA nanostructures.

## Results and discussion

### Isothermal assembly of DNA motifs and tiles in CDHP

We first explored the isothermal assembly of DNA motifs that are used commonly in DNA nanotechnology, including a double crossover (DX) motif,^37^ 3-arm junction,^38^ a 3-helix motif, ^39^ and a 3-point-star motif ^40^ (**Figure 2a** and **Figure S1**). We mixed the component strands of the DX in 1× tris-acetate-EDTA (TAE) buffer containing different concentrations of CDHP (25 to 200 mM) and incubated the samples at a constant temperature of 25 °C, 37 °C or 50 °C (**Figure S2**). We compared the assembly yields in each of these conditions to that of the samples assembled in 1× TAE with 12.5 mM Mg^2+^ using a thermal annealing protocol (**Figure 2b**). For our model nanostructure DX DNA motif, we found that assembly yields improved with both increasing temperature and CDHP concentration (**Figure S2**), with maximum assembly yield in 200 mM CDHP-containing buffer and incubated at 50 °C for 3 hours. We fixed these assembly conditions and tested the other DNA nanostructures (**Figure 2b** and **Figure S3**). Assembly yields of the 3-way junction and 3-point star were similar across the different temperatures tested while the 3-helix motif assembled with reduced efficiency, likely due to the increased complexity of the structure.

**Figure 2.**
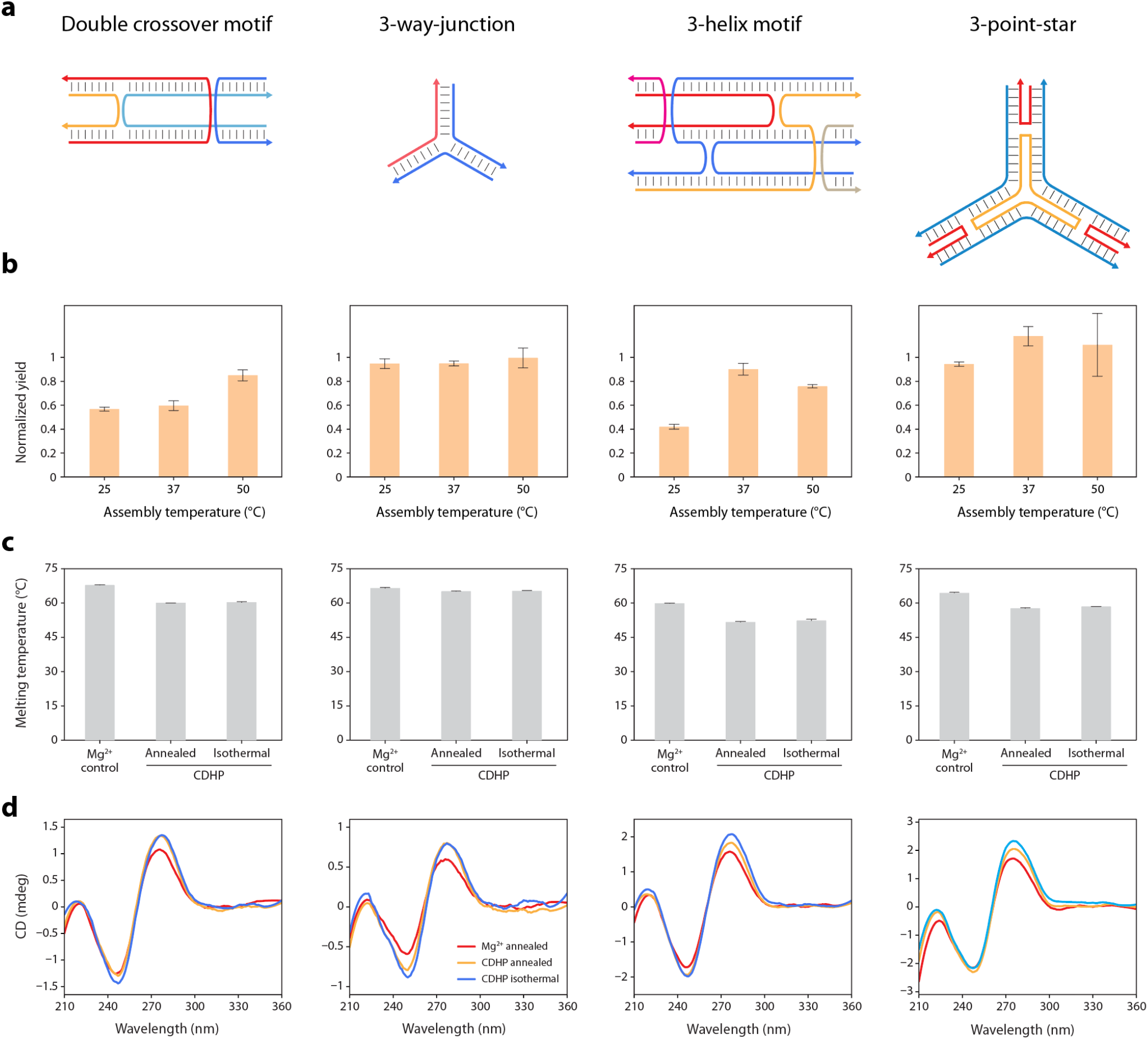
Characterization of isothermal assembly of DNA motifs in CDHP. (a) Schematic and model of DNA motifs tested here. (b) Isothermal assembly yields of each motif at different temperatures in 200 mM CDHP normalized to yields from thermal annealing in Mg^2+^. (c) Melting temperatures of motifs when assembled in different conditions. (d) CD spectra for the motifs when assembled in different conditions. Data in (b) and (c) represent mean and standard deviation from replicate experiments.

From these experiments, we established our standard assembly conditions of 1× TAE with 200 mM CDHP and isothermal incubation at 50 °C for 3 hours, which was used for further characterization studies. We looked at both the melting temperature and CD spectra of isothermally assembled structures in CDHP to determine if any changes occurred in the overall structure. There was a slight decrease in thermal stability in DX when assembled with CDHP (T_m_ ∼60 °C) compared to structures assembled in Mg^2+^ (T_m_ ∼68 °C). However, there was no change in melting temperature when comparing samples annealed or isothermally assembled in CDHP, indicating that the structures formed via isothermal assembly show same thermal stability as those assembled via thermal annealing (**Figure 2c**). We observed similar reductions in melting temperature of ∼5-10 °C for the 3-helix motif and the 3-point-star while the melting temperature of the 3-arm junction did not differ between Mg^2+^ and CDHP solutions. In the CD spectra, there were minor changes compared to the control, however, the overall spectra indicated that the structure is still B-DNA and CDHP did not adversely affect assembly (**Figure 2d**).

### Molecular dynamics simulation analysis of CDHP effect on DNA nanostructure stability

To understand the effect of CDHP on the structure of the DX motif, we performed molecular dynamics simulations (MDS) of the DX motif at 10 mM and 100 mM choline ion concentrations (**Figure 3**). We used the DX structure we previously reported^28^ and performed MDS at 300 K for three independent replicates per concentration, for 100 ns each. The dominant conformation of the motif during the simulations for 10 mM and 100 mM choline are shown in **Figure 3a**. These represent the most prevalent conformation, identified as the centroid structure of the largest cluster, by clustering the simulation snapshots based on the root mean square deviation (RMSD). The structures show that in 100 mM choline the DNA motif maintains a conformation closer to the initial ordered structure compared to the 10 mM choline simulations. This trend is also observed in the RMSD analysis of the simulation trajectories (**Figure 3b**). The stability of the motif in 100 mM choline was comparable to what we observed with 10 mM Mg^2+^. In both conditions, the RMSD reached a stable plateau of 8-10 Å after the initial equilibration period, indicating maintenance of the overall structural integrity throughout the simulations. In contrast, the 10 mM choline condition showed reduced stability with RMSD values frequently exceeding 12-15 Å and displaying greater fluctuations over time. These results show that choline can stabilize the DX structure to a degree like Mg^2+^, when present at sufficiently high concentrations, which is in alignment with experimental observations.

**Figure 3.**
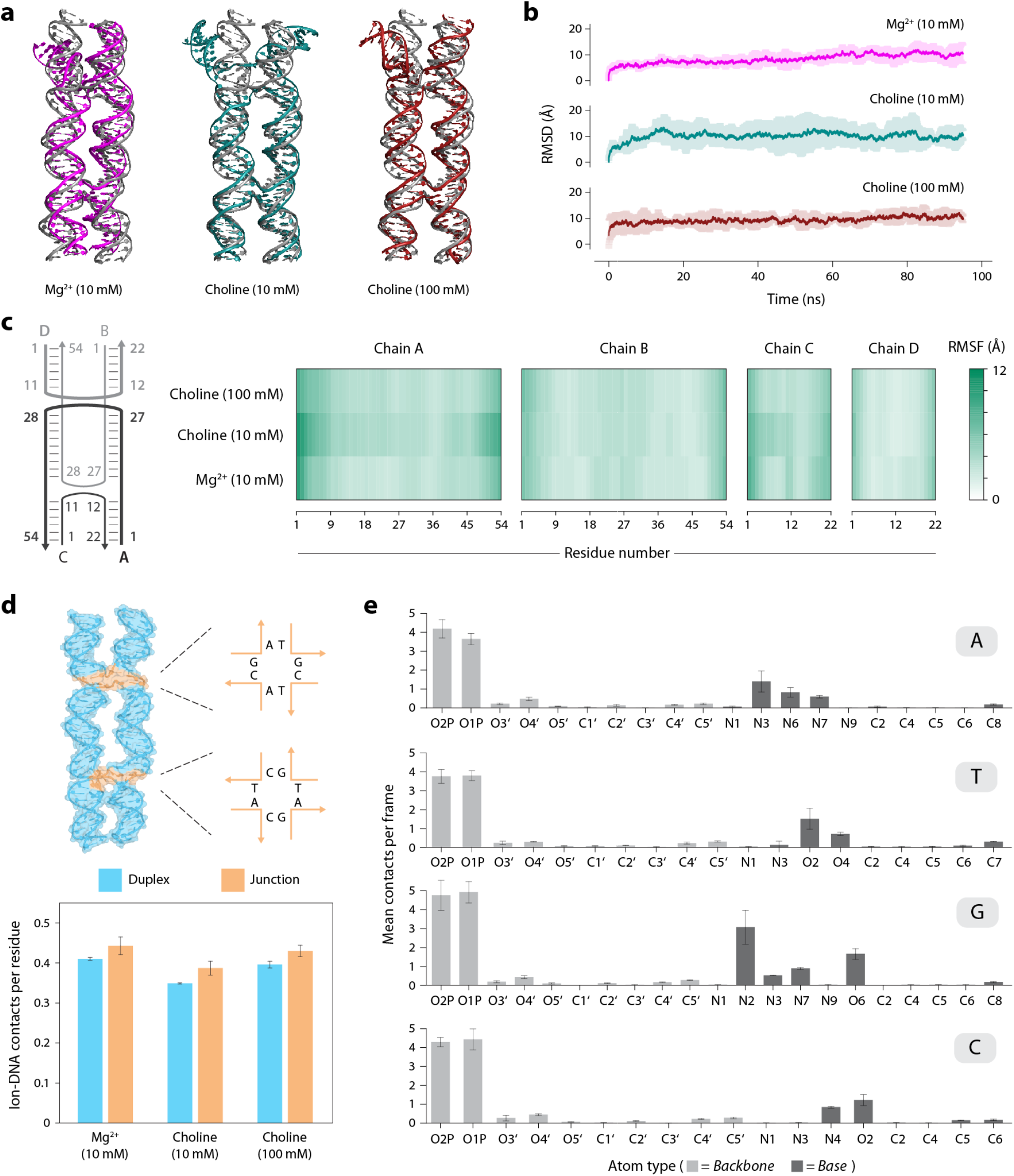
MD simulation analysis of the DX motif in CDHP. (a) Dominant conformation of the DX motif in 10 mM Mg^2+^, 10 mM choline and 100 mM choline. (b) Root mean square deviation (RMSD) of the DX motif during simulations in 10 mM Mg^2+^, 10 mM choline and 100 mM choline. (c) Root mean square fluctuation (RMSF) of nucleotides for each chain in the DX motif. (d) Interaction of choline with different structural regions of the DX motif. (e) Atom-specific contacts of the oxygen in choline with heavy atoms of the DNA obtained from simulations in 100 mM choline.

We next examined the residue-level dynamics by calculating the root mean square fluctuation (RMSF) for the nucleotides in each of the four DNA strands (chains A-D) of the motif (**Figure 3c**). The terminal residues exhibited the highest flexibility under all conditions, reflecting the lower structural constraints at the ends of the motif relative to the core. Distinct differences emerged among the individual strands. While chains B and D remained relatively rigid across all conditions, exhibiting uniformly low RMSF values along most of their lengths, chains A and C showed increased flexibility in the 10 mM choline with higher RMSF values extending inward from both the termini, compared to the 100 mM choline and 10 mM Mg^2+^ conditions. This trend is consistent with the RMSD analysis and further supports the observation that 100 mM CDHP more effectively recapitulates the stabilizing effects of Mg^2+^ on the DX motif.

To understand how the structure of the DNA affects its interactions with choline, we checked for dominant interactions between choline and the DNA by considering the junction and duplex regions of the structure separately. Choline exhibited a clear preference for the junction regions over the duplex, with the junctions showing the highest contact frequency compared to the duplex regions (**Figure 3d**). This enrichment is consistent with the increased exposure of interaction sites within the junctions, which likely makes backbone and base atoms more accessible for choline interactions. Analysis of atom-specific contacts revealed that interactions are dominated by the phosphate oxygens (O1P and O2P), indicating that electrostatic association with the negatively charged phosphate backbone is the primary binding mode (**Figure 3e**). Minor contributions from nucleobase atoms were also observed, particularly guanine N7 and O7 and to a lesser extent, cytosine N4, suggesting occasional groove-associated contacts with a slight preference for guanine-rich environments.

### Isothermal assembly of DNA origami and designer DNA crystals in CDHP

Next, we tested the effect of CDHP in the isothermal assembly of more complex structures using the DNA origami rectangle as a model system (**Figure 4**). We first characterized origami assembly using non-denaturing agarose gel electrophoresis (AGE) and observed a band corresponding to the origami rectangle in all tested isothermal conditions, with increasing assembly efficiency as temperature increases (**Figure S4**). To validate origami structural formation, we performed atomic force microscopy (AFM) imaging that confirmed successful formation of the expected origami structure at 37 °C and 50 °C. Isothermal assembly was inefficient at 25 °C as only a few partially assembled rectangular structures were observed by AFM imaging.

**Figure 4.**
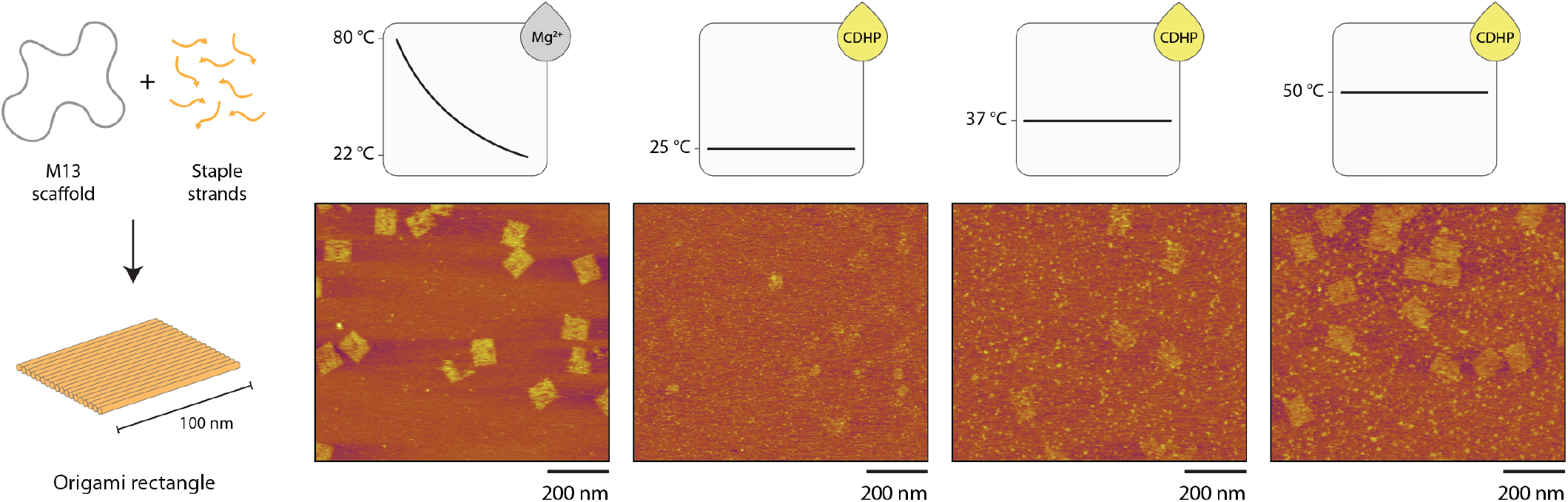
Isothermal assembly of DNA origami in CDHP. Schematic showing DNA origami self-assembly (left) and AFM images of the DNA origami rectangle assembled by a thermal annealing protocol in buffer containing Mg^2+^ or isothermal assembly in buffer containing CDHP at 25, 37 or 50 °C.

Based on the successful assembly of finite nanostructures in CDHP, we then explored the isothermal assembly of hierarchical DNA lattices using CDHP. Designer DNA crystals based on the tensegrity triangle DNA motif have been constructed before, where rational design of the motif allows the creation of 3D lattices with pre-defined cavities. Here, we used a tensegrity triangle that contains three double helical edges with 31 bp each (corresponding to 3 helical turns), connected at the vertices by DNA crossovers (**Figure 5a** and **Figure S5**). The edges are tailed by 2-nt sticky ends that allow the individual motifs to self-assemble into a 3D crystal. In contrast to typical thermal annealing in buffer containing Mg^2+^, here we tested the self-assembly of these motifs into 3D crystals in 200 mM CDHP containing buffer (**Figure S6**). Motifs were isothermally assembled in 25, 37 or 50 °C and set up for crystallization using the hanging drop vapor diffusion method (**Figure 5b**). We observed crystals in all three conditions, with the typical crystal habit of rhombohedral shape observed for these crystals in previous studies^41–43^ (**Figure 5c**).

**Figure 5.**
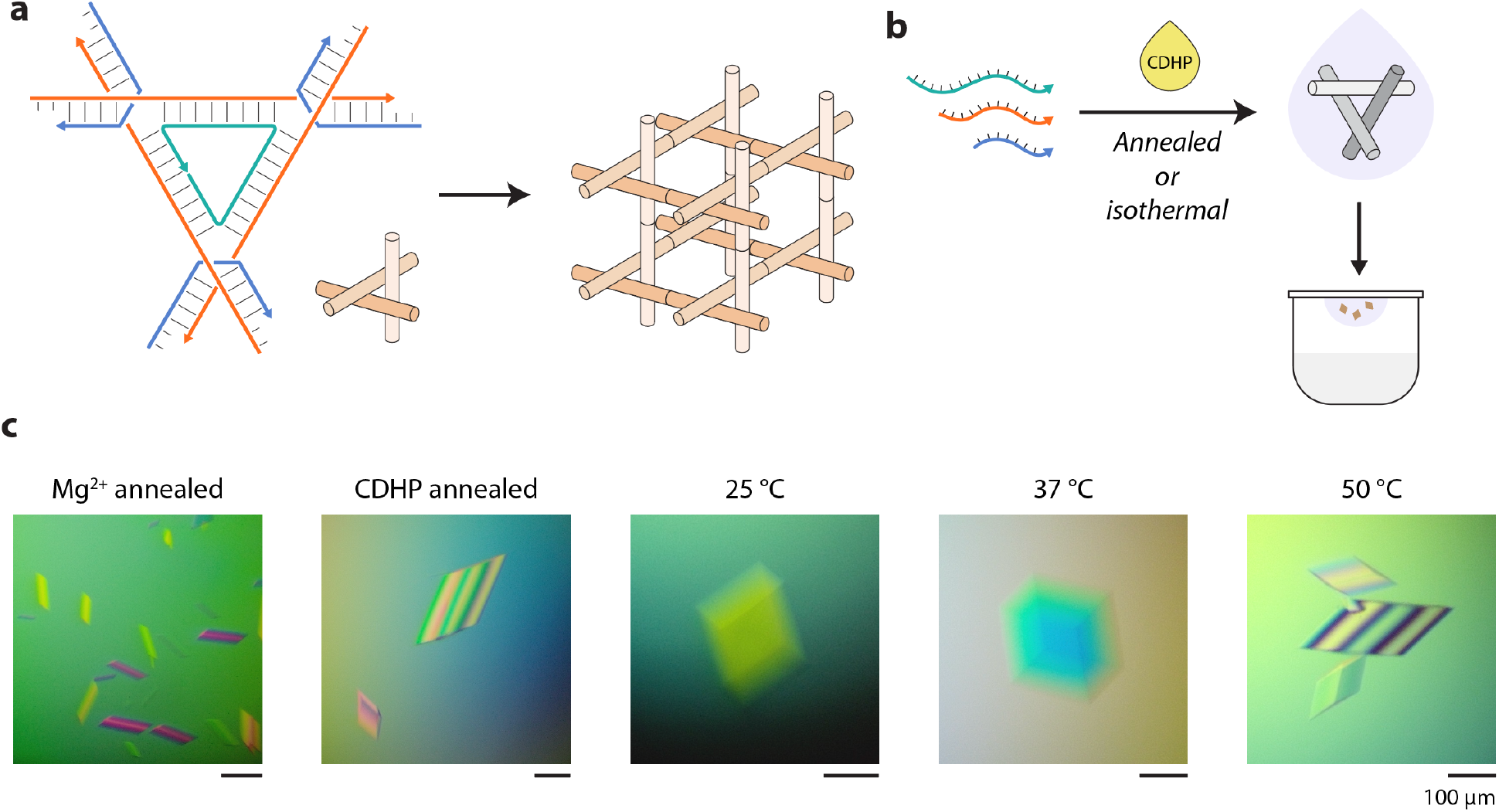
3D DNA crystals from isothermally assembled tensegrity triangle DNA motifs. (a) Schematic of the tensegrity triangle motif containing 3 helical turns per edge. The motif assembles into a 3D lattice via sticky end cohesion. (b) Isothermally assembled motifs were set up for crystallization using the hanging drop vapor diffusion method. (c) Crystals obtained from motif isothermally assembled at 25, 37 or 50 °C.

### Enhanced biostability and biological characteristics of CDHP-assembled structures

Previous studies have shown that the addition of CDHP in the assembly process of functional nucleic acids significantly increased the half-time of nucleic acids when incubated with nucleases.^24^ This effect was also seen in our recent work when annealing DNA nanostructures with CDHP.^25^ To determine if this effect was disrupted by the isothermal assembly process, we incubated isothermally assembled DX with different nucleases and used the DX annealed in Mg^2+^ as the control structure. More specifically, the DX structures were incubated for 30 minutes with a range of concentrations of T5 exonuclease, exonuclease V, and exonuclease III (**Figure 6a-c** and **Figure S7**). In all tested conditions the DX isothermally assembled with CDHP showed higher nuclease resistance compared to the control structure annealed in Mg^2+^. In T5 exonuclease, the Mg^2+^-assembled structure was almost fully degraded in 0.5 U while the CDHP-assembled structure was degraded with 1 U enzyme. In enzymes exonuclease V and exonuclease III, the Mg^2+^-assembled structures were 60% and ∼95% degraded at the highest nuclease concentration used herein, while the CDHP-assembled structures were almost fully intact. For potential biological applications, we tested cell viability and internalization of DNA nanostructures. We observed no change in cell viability of HepG2 cells incubated with DNA nanostructures isothermally assembled in CDHP when compared to untreated cells or Mg^2+^-assembled structures (**Figure 6d**). Further, isothermal assembly in CDHP did not affect cell internalization of DNA nanostructures (**Figure 6e-f**). These results further enhance our prior work that showed enhanced cell receptor recognition of aptamer-functionalized DNA nanostructures annealed and presented in a CDHP containing buffer.^25^

**Figure 6.**
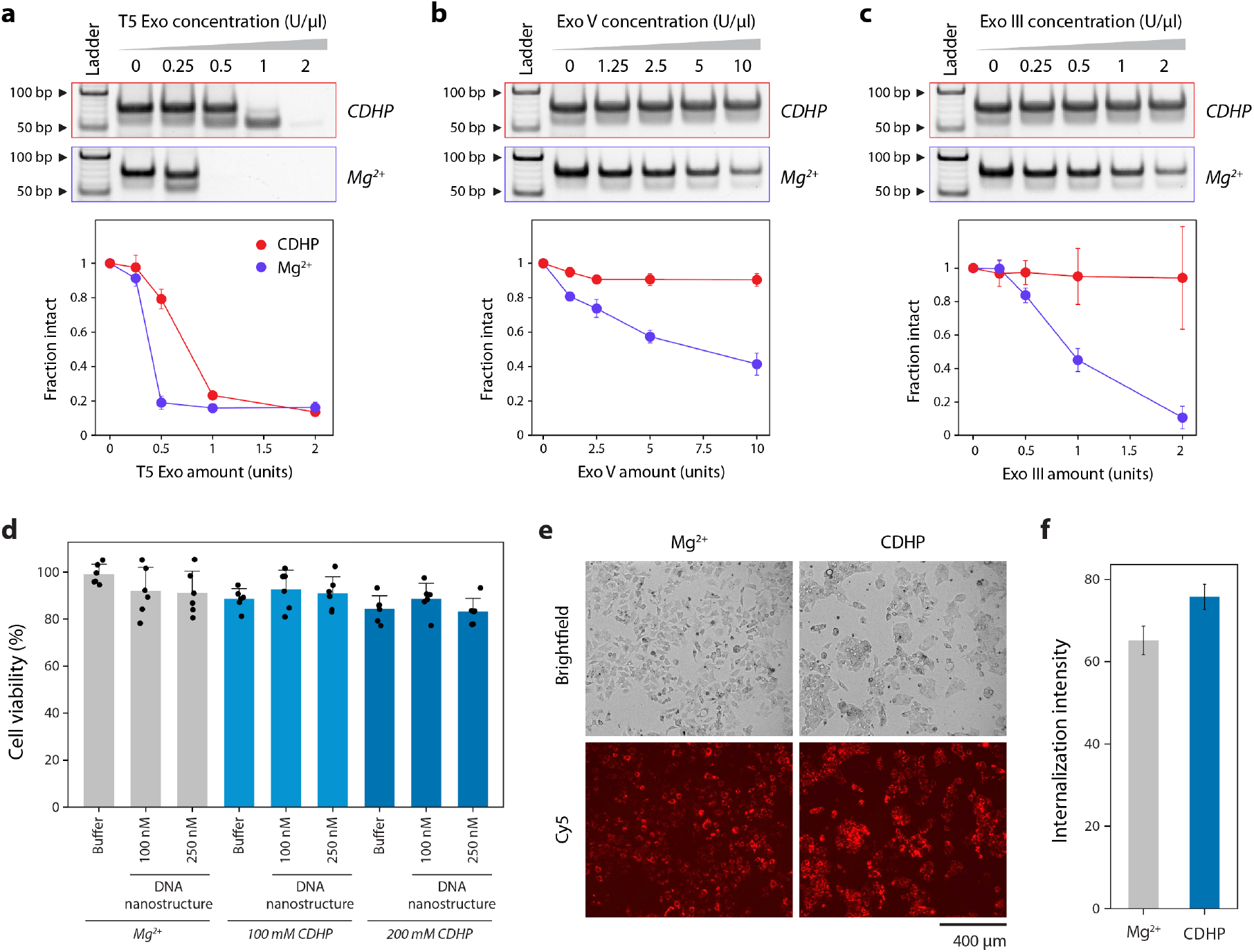
Towards biological feasibility. Enzymatic degradation of DX motif isothermally assembled with CDHP in (a) T5 exonuclease, (b) Exo V and (c) Exo III nuclease. Data in a-c represent mean and error propagated from SDs of experiments performed in triplicates. (d) Cell viability of HepG2 cells incubated with DNA nanostructure annealed in Mg^2+^ or isothermal assembled in CDHP. Data represents mean and error propagated from SDs of six biological replicates. (e-f) Internalization of a model DNA nanostructure in HepG2 cells. Data in (a), (b) and (c) represent mean and standard deviation from replicate experiments. Data in (d) and (f) were obtained from cell experiments performed with six biological replicates.

### Mechanistic insights into choline-mediated enhanced nuclease resistance of DNA nanostructures

Since all atom MDS can provide atomic level resolution on ion interactions with DNA, we compared the interactions of choline with the DX motif relative to Mg^2+^ to understand the difference in nuclease resistance observed in the two cases. Previous studies have shown that choline-containing ionic liquids stabilize nucleic acids through direct and multimodal interactions with DNA, including groove and backbone binding.^44^ Choline has been reported to form persistent hydrogen-bonding networks with DNA and to preferentially localize in accessible groove environments, distinguishing its behavior from that of hydrated metal cations.^45,46^ Furthermore, the “cage-effect” of CDHP ionic liquids has been shown to suppress hydrolysis and nuclease-mediated degradation by limiting water accessibility while preserving nucleic acid structure and function.^24^ We calculated the Solvent Accessible Surface Area (SASA) of the DNA from the MDS trajectories. Choline ions reduce the SASA for the DAO motif compared to the Mg^2+^ (**Figure 7a**). DNA proximity distributions, calculated as the distribution of the distance between the oxygen of the choline (or the Mg^2+^ ion) with the nearest heavy atoms in the DNA, highlighted distinct interaction mechanisms for choline and Mg^2+^ **(Figure 7b**). Choline displayed a broad peak at ∼ 3 Å from the DNA, consistent with the direct backbone contact, whereas Mg^2+^ showed a narrower peak centered around ∼4 Å, indicative of a more ordered, water-mediated interaction. Representative structures from the simulations show that Mg^2+^ typically retains its first hydration shell and interacts with the DNA through bridging water molecules, while choline most often contacts the backbone directly (**Figure 7c-e**). While some of these interactions might be a result of the difference in size of the choline ions compared to the Mg^2+^, the water mediated interactions in Mg^2+^ suggest that the ions remain further away from the DNA compared to choline allowing water to easily access the DNA surface. Together, these results show that although high concentrations of choline can provide structural stabilization comparable to Mg^2+^, the underlying mechanisms differ: Mg^2+^ stabilized DNA through a slightly longer water-mediated electrostatic interactions, where as choline stabilizes through direct close-range contacts concentrated at the junctions. These properties provide a plausible explanation for the enhanced DNA integrity observed in choline-containing systems relative to conventional Mg^2+^ stabilized environments.

## Conclusion

The results presented here demonstrate that the conditions required for successful DNA nanostructure isothermal assembly are not limited to using solutions containing metallic counter ions. Our study shows that DNA nanostructures of increasing complexity can be assembled isothermally in CDHP. Further, this is the first report of designer DNA crystal assembly in hydrated ionic liquids. Enhanced enzymatic resistance associated CDHP-based assembly was maintained when the structure is assembled isothermally. Isothermal assembly in CDHP also does not impact the shape or thermal stability of the tested DNA nanostructures, further supporting the use of hydrated ionic liquids such as CDHP as an alternative to the traditionally used Mg^2+^, with the added benefits of enhanced biostability and functionality.^25^ Isothermal assembly also allows for a much simpler assembly process, and reduced assembly temperatures would allow for single pot synthesis with guest molecules. While a previous report showed assembly of DNA nanostructures in a deep eutectic solvent glycholine, such isothermal assembly involved an initial heating step at 70 °C.^22^ In contrast, we show assembly in an entirely isothermal process at temperatures at or below 50 °C. Our MD simulations provide molecular level insight into the interactions between CDHP-derived choline and DNA nanostructure, highlighting that choline can stabilize the structure comparably to Mg^2+^ while utilizing a direct and close-range interaction in contrast to the predominantly water-mediated interactions of Mg^2+^. This provides a plausible explanation for the enhanced nuclease resistance observed in CDHP-containing environments. The simulations further suggest that choline preferentially associates with the structurally open junction regions and to a lesser extent, guanine-rich sites. Together these findings offer design principles for the rational optimization and tuning of assembly conditions for a variety of other DNA nanostructures. In addition to improving biological compatibility of DNA nanostructures, this strategy could be amenable to other utilities such as dynamic control stimulated by light or nucleic acids.

**Figure 7.**
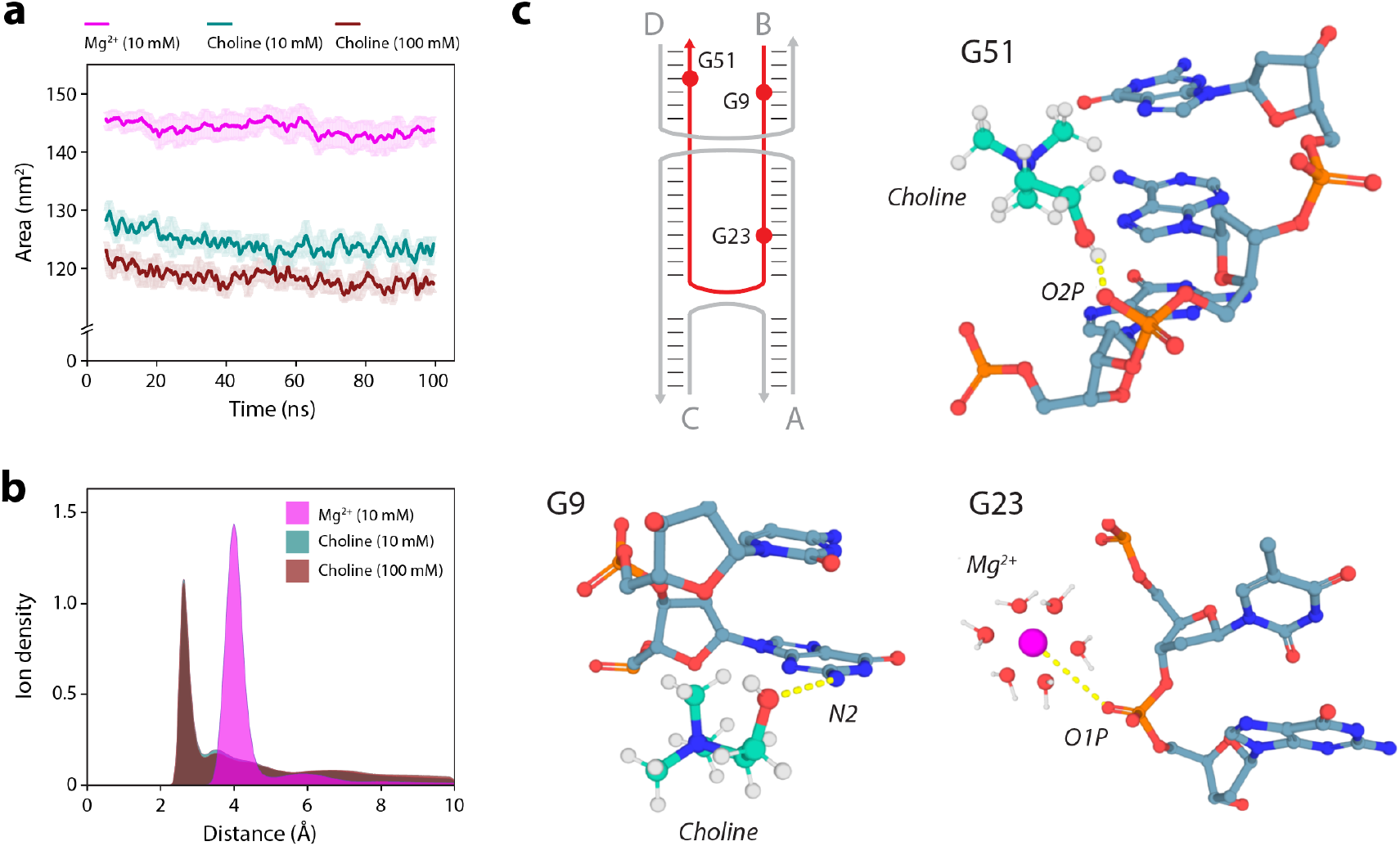
*Interaction of choline and Mg*^*2+*^ *with the DX motif*. (a) Solvent accessible surface area (SASA) of the ion bound DX motif in the presence of choline and Mg^2+^. (b) Distance distribution of the choline and Mg^2+^ ions relative to the DNA. Representative MD snapshots illustrating interaction modes of (c) choline with DNA backbone, (d) choline with nucleobase, and (e) Mg^2+^ with DNA (water mediated).

## Associated content

Materials and methods, additional experimental results and sequences used.

## Author contributions

The project was supervised by A.R.C and X.W. Experiments were designed by H.T., L.N., D.G., X.W. and A.R.C. DNA motif assembly and characterization was performed by H.T. and G.G. Biostability assays and crystal assembly were performed by H.T. DNA origami assembly and characterization was performed by L.N. and D.G. MD simulations were performed by Z.N. and K.M. with supervision from S.V. Cell studies were performed by C.M. under the supervision of T.J.B. Figures were made by H.T. and A.R.C. The paper was written by H.T. and A.R.C. with edits from D.G., S.V. and X.W. All authors provided feedback on the manuscript.

## Acknowledgments

Research reported in this publication was supported by the National Institutes of Health (NIH) through National Institute of General Medical Sciences (NIGMS) under award number R35GM150672 to A.R.C. and through the National Institute of Environmental Health Sciences (NIEHS) under award number R01ES026856 to T.J.B. This work was also supported in part by grant from NSF 2127436 to X.W. H.T. was supported by the T32 Training Grant awarded to The RNA Institute from the NIH through NIGMS under award number T32GM132066, “RNA Science and Technology in Health and Disease”. This manuscript is the result of funding in whole or in part by the National Institutes of Health (NIH). It is subject to the NIH Public Access Policy. Through acceptance of this federal funding, NIH has been given a right to make this manuscript publicly available in PubMed Central upon the Official Date of Publication, as defined by NIH.

