## Supporting information for "Multi-dimensional DNA nanostructures isothermally assembled in hydrated ionic liquids"

**Preparation of DNA complexes**

*DNA motifs.* Strands were purchased from Integrated DNA Technologies with standard desalting. Strands were combined in Tris-acetate-EDTA buffer containing 40 mM Tris base (pH 8), 20 mM acetic acid, and 2mM EDTA (1× TAE). The buffer in control samples also contained 12.5 mM $Mg^{2+}$ (1× TAE-$Mg^{2+}$) and the buffer for CDHP-containing samples contained 200 mM CDHP adjusted to pH 7 using NaOH (1× TAE-CDHP) unless otherwise stated. Strands for DAO, 3WJ, and TX where combined in equimolar ratios and 3PS combined the strands in a 3:3:1 ratio. Annealed samples were assembled using a step-wise annealing protocol on a thermocycler using these steps: 90°C for 3 min, 65°C for 20 min, 45°C for 20 min, 37°C for 30 min, 20°C for 30 min, then cooled to 4°C. Isothermal samples were incubated at indicated temperatures for 3 hours. Once it was established that 50°C incubation produced the best yields, it was used for all characterization studies.

*DNA origami.* Rectangular DNA origami nanostructures were assembled by mixing the M13mp18 scaffold strand with the respective staple strands at a 1:5 molar ratio with a final DNA origami concentration of 20 nM in 1× TAE buffer (40 mM Tris, 20 mM acetic acid, and 1 mM EDTA) containing 125 mM CDHP (pH 7.4, adjusted using NaOH).For thermally annealed samples, the mixture was gradually cooled from 80 °C to 22 °C over 16 hours in a Biometra TRIO Thermal Cycler. For isothermal assembly, the mixtures were incubated at different constant temperatures (25, 37 and 50 °C) for 16 hours.

*DNA crystals.* Crystals were grown in 6 µl hanging drops containing 2.5 µM DNA tensegrity triangle solution annealed or isothermally assembled in 1× TAE containing 12.5 mM $Mg^{2+}$ or 200 mM CDHP. DNA samples in hanging drops contained with 0.825 M ammonium sulfate and drops were equilibrated against a 600 ml reservoir of 1.75 M ammonium sulfate at 20 °C. The following protocol was used for annealed samples: 90 °C for 3 minutes, 65 °C for 20 minutes, 45 °C for 20 minutes, 37 °C for 30 minutes, 20 °C for 30 minutes. For isothermal assembly, samples were incubated at 25, 37 or 50 °C for 3 h. Crystals were imaged using a Zeiss SteREO Discovery V12 microscope.

**Gel electrophoresis**

*Polyacrylamide gel electrophoresis.* Non-denaturing gels were prepared using 19:1 acrylamide:bis-acrylamide solution (AccuGel, National Diagnostics). Gels were run using 1× TAE with 12.5 mM $Mg^{2+}$ running buffer at a constant volage in 4°C. Gels were stained in 0.5× GelRed (Biotium) solution. Gels were imaged using Gel Doc XR+ (BioRad) with standard settings for GelRed illumination and analyzed using ImageLab software.

*Agarose gel electrophoresis.* Gel analysis of the isothermally assembled DNA origami structures was performed on 1% agarose gels with 1× TAE as running buffer. SYBR Green (Thermo Fisher Scientific) was incorporated into the gels prior to casting. Samples were separated at 7.5 V/cm for 2 h on ice. Gel images were acquired using a UVP GelStudio imaging platform.

### Circular dichroism

DNA motifs were prepared at 1 µM concentration in 1× TAE-$Mg^{2+}$ or 1× TAE-CDHP. Control samples were annealed using the standard thermal melting protocol. Isothermal samples were incubated at 50 °C for 3 hours. CD measurements were taken immediately following assembly. A 1 mm quartz cuvette (Hellma Analytics) was used. Measurements were taken using a Jasco-815 CD spectrometer while at room temperature. Measurements were taken every 1 nm from 200 nm to 360 nm at a scanning rate of 50 nm/min. Measurements are an average of 3 accumulations.

### UV melting

DNA motifs were prepared at 1 µM concentration in 1× TAE-$Mg^{2+}$ or 1× TAE-CDHP and analyzed immediately following their assembly. Measurements were taken using a Cary 3500 UV-Visible Spectrophotometer (Agilent). Absorbance at 260 nm was recorded from 15°C to 95°C every 0.5°C. Melting curves were normalized 0-1 and fitted to the Boltzman curve using OriginLab software. Melting temperatures were obtained using the first derivative of the Boltzman curve.

### Molecular dynamics (MD) simulations

The initial all-atom structure of the DAO motif was modeled in Molecular Operating Environment (MOE)[1] and reported in our previous work.[2] Briefly, two double stranded DNAs were generated, and strand breaks and joins were introduced at the appropriate locations to create the double crossover motif. MD simulations were then performed using GROMACS 2019.4[3] to assess the stability and interaction of the structure with choline ions. The amber99-bsc01 force field was used for DNA.[4] The choline cation was parameterized using HF/6-31G* electrostatic potentials calculated in Q-Chem. Atomic charges were derived using the RESP method implemented in Antechamber following the AMBER force-field protocol,[5–7] and the resulting parameters were converted for use in GROMACS. All simulations were carried out at 300 K for 10 mM and 100mM choline ion concentration. Three replicate 100 ns simulations were performed for each concentration.

The MDS incorporated a leap-frog algorithm with a 2-fs time step to integrate the equations of motion. The system was simulated using the velocity rescaling thermostat to maintain the temperature. [8] The pressure was maintained at 1 atm using the Berendsen barostat for equilibration.[9] Long-range electrostatic interactions were calculated using the particle mesh Ewald (PME) algorithm with a real-space cut-off of 1.0 nm.[10] Lennard–Jones interactions were truncated at 1.0 nm. Water molecules were represented with the TIP3P model,[11] and the LINCS algorithm[12] was used to constrain the motion of hydrogen atoms bonded to heavy atoms. The system was subjected to energy minimization to prevent overlap of atoms, followed by a 100 ns molecular dynamics simulation. The first 5 ns of the trajectory were excluded from the analysis for system equilibration. Coordinates were stored every 2 ps for further analysis. The simulations were visualized using PyMOL.[13] The Root Mean Square Deviation (RMSD) and Root Mean Square Fluctuations (rmsf) were calculated and analyzed using the GROMACS tools rms and rmsf respectively.[3]

The DNA motif was divided into junction 1, junction 2 and duplex regions for ion contact analysis. Junctions were defined as four nearest base pairs at each of the crossover regions of the DAO motif. The mindist tool in GROMACS[3] was used to calculate the distance, number and type of contacts between choline ions and the DNA. Only heavy atoms of the DNA were used in the calculations. All hydrogens were excluded. Two atoms were considered in contact if the distance between them was less than 3.5 Å. The results were normalized by the number of nucleotides. Solvent-accessible surface area (SASA) values were calculated using the GROMACS tool, gmx sasa, with the analysis performed on the DNA in the presence of ions.

**AFM imaging**

The mica substrates were rinsed with double-distilled water and dried under compressed air prior to sample deposition. Isothermally assembled DNA origami samples were diluted to a final concentration of 2–3 nM in 1× TAE buffer and 5 µL of the diluted sample was deposited onto the mica surface. After a 5 min incubation, the surface was gently washed three times with deionized water to remove excess material, followed by the addition of 40 µL deionized water for imaging under buffer conditions. The imaging was performed in tapping mode using a fluid cell on a Multimode Asylum Research Cypher S AFM equipped with an MSNL-10 probe (Bruker Nano Inc.). The acquired AFM images were processed using Gwyddion (v2.6.1), where background flattening was applied to accurately visualize the height profiles of the DNA origami structures.

**Nuclease degradation assays**

DX samples were prepared at 0.25 µM in either 1× TAE-$Mg^{2+}$ or 1× TAE-CDHP. Samples containing $Mg^{2+}$ were assembled using the standard annealing protocol and samples containing CDHP were assembled isothermally at 50°C for 3 hours. Samples were combined with 10× NEBuffer 1 for a 1× NEBuffer 1 final concentration. 9 µl of each DX sample with NEBuffer 1 were combined with 1 µl of each enzyme at varying concentrations. This was then incubated at 37 °C for 30 minutes.

**Cell viability assay**

Cell viability was assessed using the MTT (3-(4,5-dimethylthiazol-2-yl)-2,5-diphenyltetrazolium bromide) assay, which measures metabolic activity of viable cells. HepG2 Cells were seeded in 96-well plates at a density of $1.5 \times 10^4$ cells per well and allowed to attach overnight under standard culture conditions (37 °C, 5% $CO_2$). The next day, cells were treated with DNA tensegrity triangle motifs assembled in CDHP or $Mg^{2+}$ and their respective 1× TAE buffers as controls for 24 hours. Following treatment, MTT reagent was added to each well at a final concentration of 0.5 mg/ml and incubated for 3–4 hours at 37 °C to allow viable cells to reduce MTT into insoluble purple formazan crystals. After incubation, the culture medium was carefully removed and 100 µl of dimethyl sulfoxide (DMSO) was added to each well to dissolve the formazan crystals. Plates were gently agitated for 10–15 minutes to ensure complete solubilization. Absorbance was measured at 570 nm and cell viability was calculated as the percentage of absorbance relative to the untreated control group (n = 6 biological replicates).

RM One-Way ANOVA analysis for cell viability shown that the p-values are non-significant when compared to buffer-only controls.

| | | |
|---|---|---|
| 100 nM DNA nanostructure in 12.5 mM $Mg^{2+}$ | ns | 0.1521 |
| 250 nM nanostructure in 12.5 mM $Mg^{2+}$ | ns | 0.0968 |
| 100 nM DNA nanostructure in 100mM CDHP | ns | 0.3239 |
| 250 nM DNA nanostructure in 100 mM CDHP | ns | 0.5447 |
| 100 nM DNA nanostructure in 200mM CDHP | ns | 0.2177 |
| 250 nM DNA nanostructure in 200mM CDHP | ns | 0.8712 |

**DNA structure internalization assay**

Cellular uptake of DNA structures was assessed using Cy5-labeled DNA tensegrity triangle motifs (strand 3TS3 was modified to contain a Cy5 dye). HepG2 cells were seeded on glass-bottom dishes or coverslips in 12-well plates at a density of $1.5 \times 10^5$ cells per well and allowed to attach overnight under standard culture conditions (37 °C, 5% $CO_2$). Cy5-labeled DNA structures were prepared at 5 µM concentrations and added to the cells at 50 nm final concentration. 1× TAE buffer was used as the control. Cells were incubated for the specified time (1-6 hours) to allow internalization. Fluorescence images were acquired using an EVOS Fluorescence Microscope with the appropriate Cy5 filter at 3-hour timepoint. Quantification of internalization was performed using ImageJ. Background fluorescence was subtracted, and the Corrected Total Cell Fluorescence (CTCF) intensity and percentage of DNA internalization was calculated per condition. This experiment was conducted in triplicate.

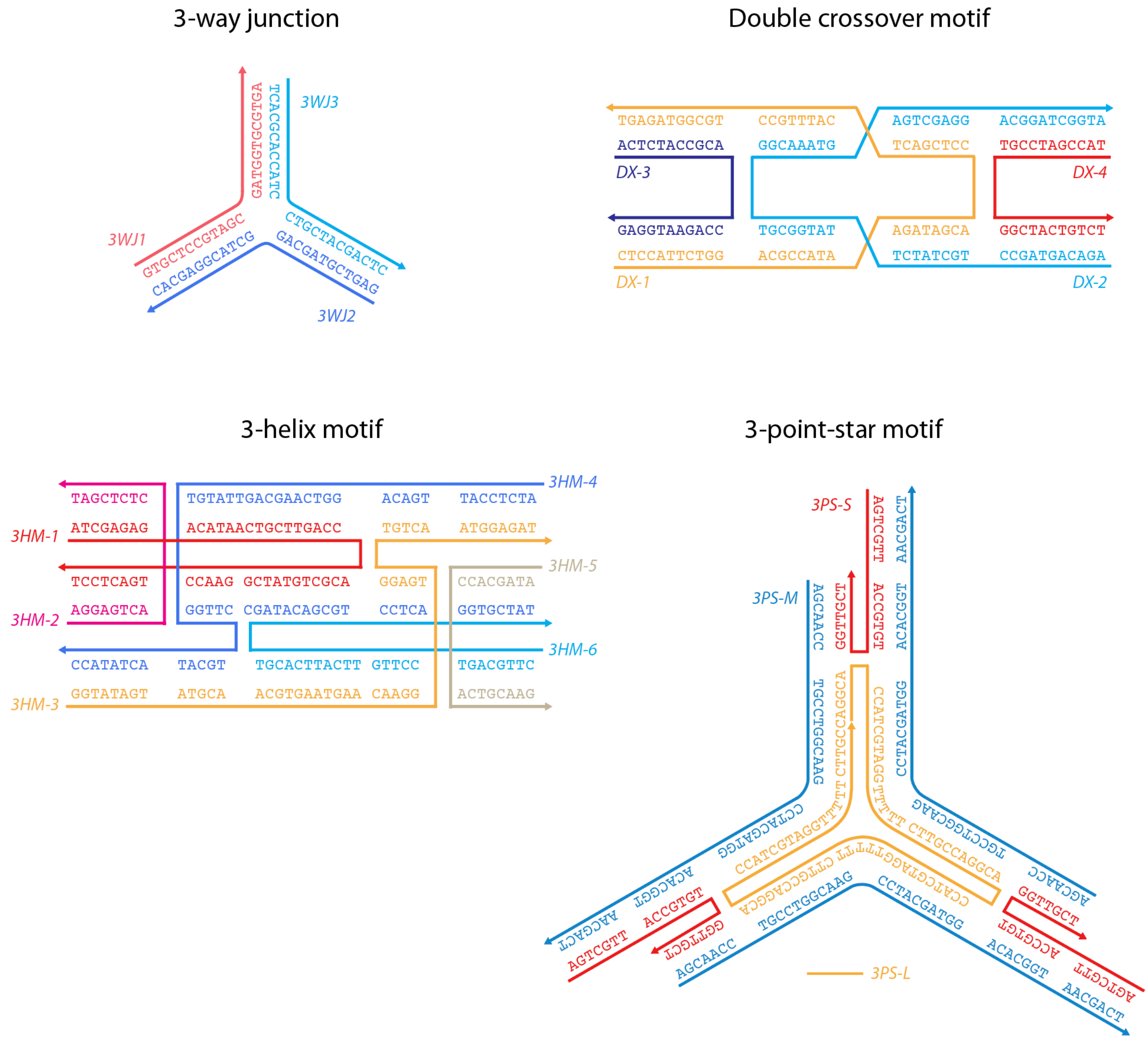


**Figure S1.** Scheme and sequences of DNA nanostructures used in the study.

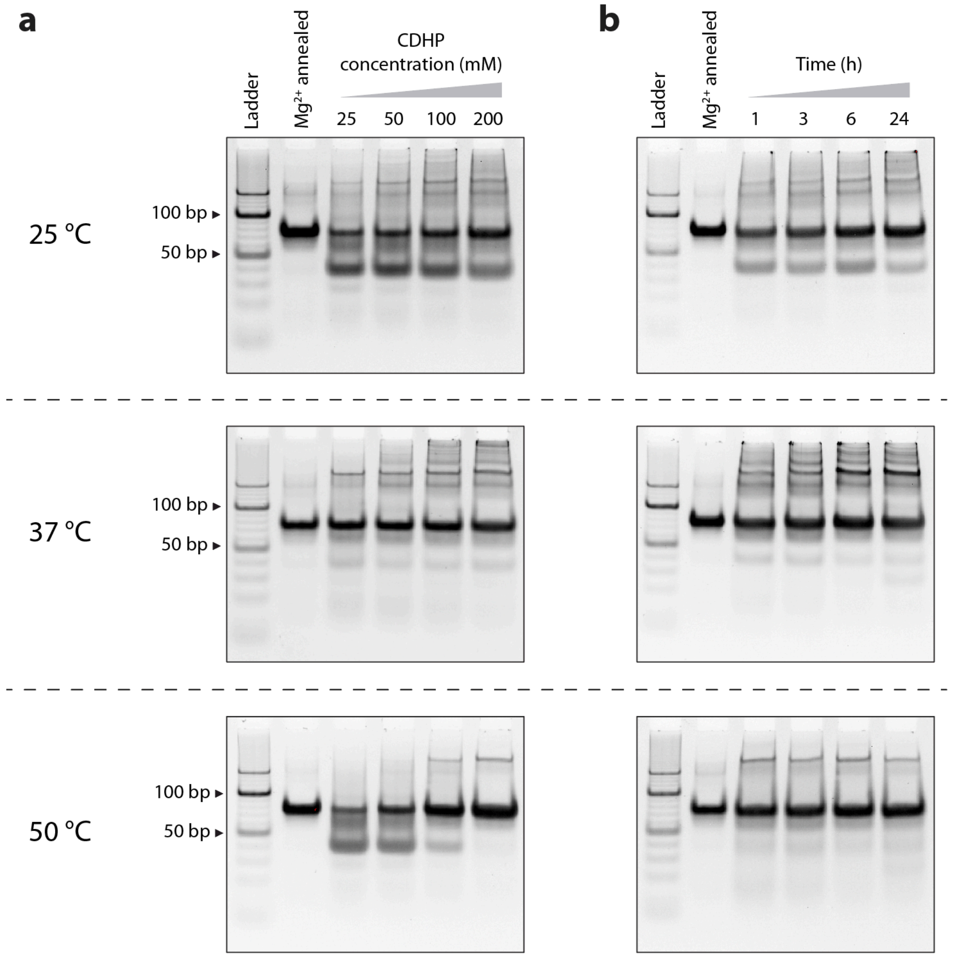


**Figure S2.** Isothermal assembly of the model nanostructure DX motif in (a) different concentrations of CDHP for 3 hours and (b) at different incubation times in 200 mM CDHP.

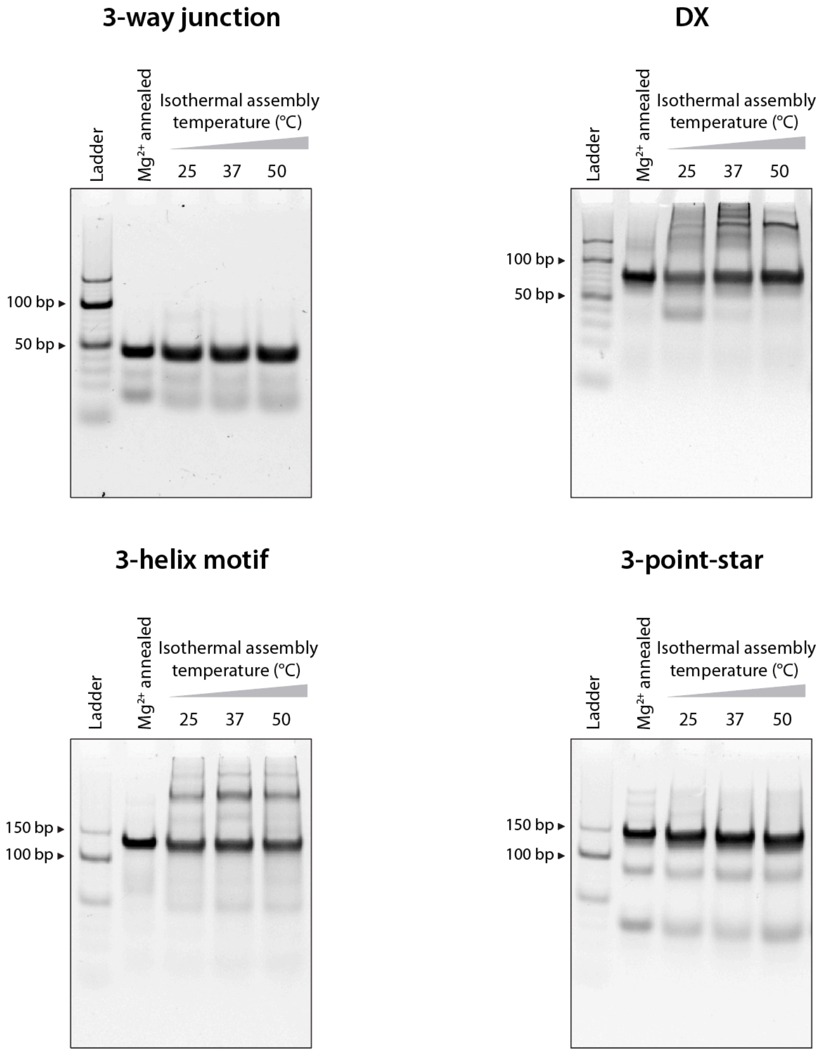


**Figure S3.** Isothermal assembly of the different DNA motifs. Gels used for analysis shown in Figure 2b.

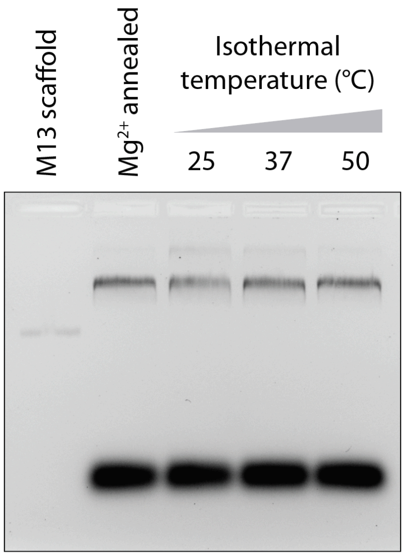


**Figure S4.** Non-denaturing agarose gel for the isothermal assembly of DNA origami rectangle.

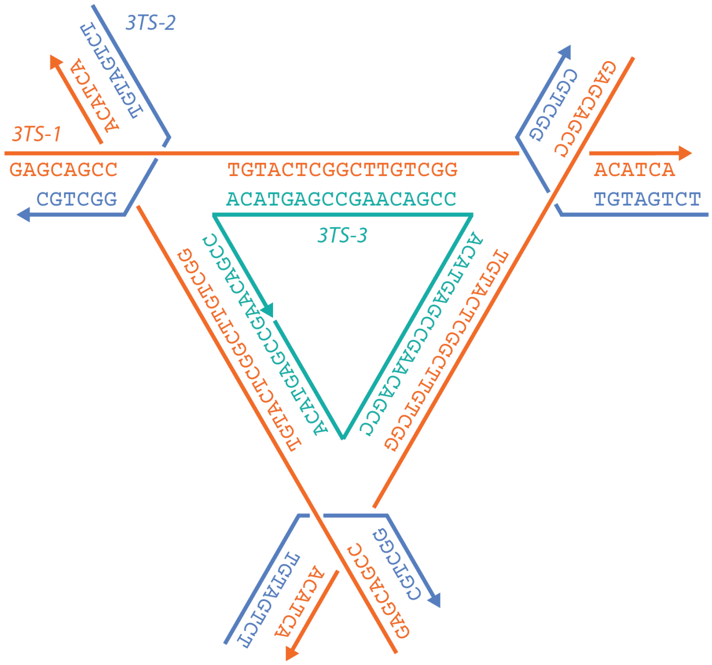


**Figure S5.** Scheme and sequences of the DNA tensegrity triangle used for self-assembled crystals.

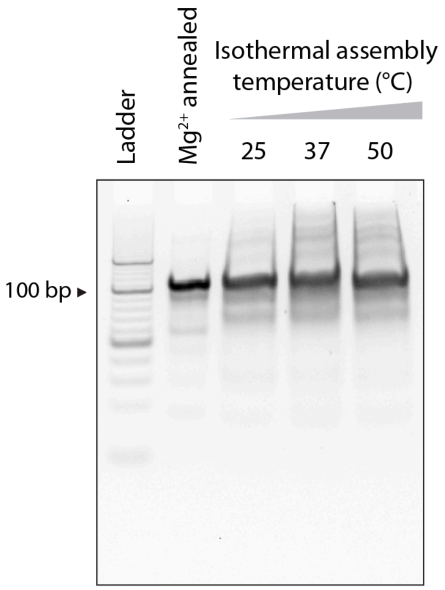


**Figure S6.** Non-denaturing gel showing the assembly of DNA tensegrity triangle.

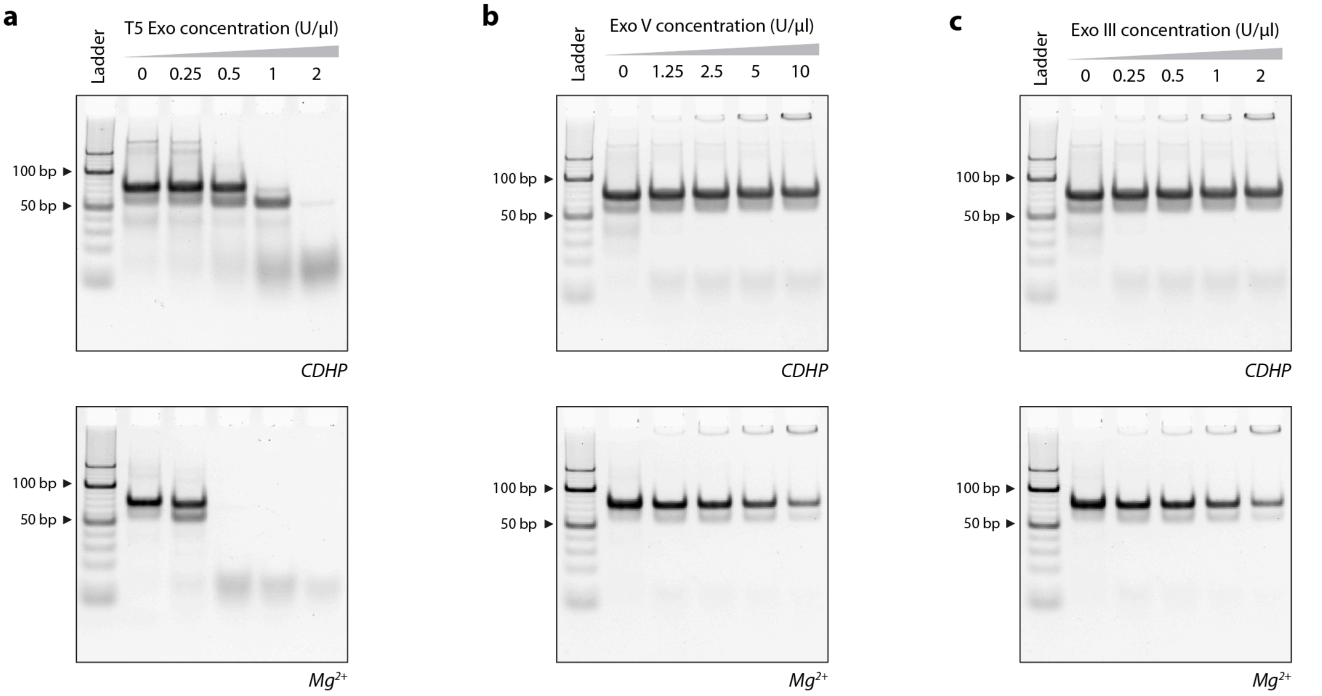


**Figure S7.** Non-denaturing gels for nuclease degradation analysis of the DX DNA motif. (a) T5 exonuclease, (b) Exo V and (c) Exo III nuclease. Full images of gels shown in Figure 6a-c.

**Table S1**. Sequences for DNA motifs used in this study (written 5′-3′)

| Strand | Sequence |
|---|---|
| **Double crossover motif** | |
| DX-1 | CTCCATTCTGGACGCCATAAGATAGCACCTCGACTCATTTGCCTGCGGTAGAGT |
| DX-2 | AGACAGTAGCCTGCTATCTTATGGCGTGGCAAATGCGTCGAGGACGGATCGGTA |
| DX-3 | ACTCTACCGCACCAGAATGGAG |
| DX-4 | TACCGATCCGTGGCTACTGTCT |
| **3-way junction** | |
| 3WJ-1 | AAGTGCTCCGTAGCGATGGTGCGTGATT |
| 3WJ-2 | AATCACGCACCATCCTGCTACGACTCAA |
| 3WJ-3 | TTGAGTCGTAGCAGGCTACGGAGCACTT |
| **3-point star** | |
| 3PS-1 | AGTCGTTACCGTGTGGTTGCT |
| 3PS-2 | AGCAACCTGCCTGGCAAGCCTACGATGGACACGGTAACGACT |
| 3PS-3 | AGGCACCATCGTAGGTTTTTCTTGCCAGGCACCATCGTAGGTTTTTCTTGCCAGG CACCATCGTAGGTTTTTCTTGCC |
| **3-helix motif** | |
| TX-1 | ATCGAGAGACATAACTGCTTGACCACGCTGTATCGGAACCTGACTCCT |
| TX-2 | AGGAGTCACTCTCGAT |
| TX-3 | GGTATAGTATGCAACGTGAATGAACAAGGTGAGGTGTCAATGGAGAT |
| TX-4 | ATCTCCATTGACAGGTCAAGCAGTTATGTGGTTCTGCATACTATTACC |
| TX-5 | ATAGCACCACTGCAAG |
| TX-6 | CTTGCAGTCCTTGTTCATTCACGTCGATACAGCGTCCTCAGGTGCTAT |
| **3-turn tensegrity triangle** | |
| 3TS-1 | GAGCAGCCTGTACTCGGCTTGTCGGACATCA |
| 3TS-2 | TCTGATGTGGCTGC |
| 3TS-3 | CCGAGTACACCGACAAGCCGAGTACACCGACAAGCCGAGTACACCGACAAG |

**Table S2.** Staples used for assembly of DNA rectangle origami.

| Sequence |
|---|
| TGCCTTTAGTCAGACGATTGGCCTGCCAGAATAAGCTGCTCTCCAAA |
| TGAACAAACAGTATGTTAGCAAACTAAAAGAAAAGCTGCTCTCCAAA |
| CCAGACGAGCGCCCAATAGCAAGCAAGAACGCAAGCTGCTCTCCAAA |
| TTGAATTATGCTGATGCAAATCCACAAATATAAAGCTGCTCTCCAAA |
| ATCAACAGTCATCATATTCCTGATTGATTGTTAAGCTGCTCTCCAAA |
| TCACAATCGTAGCACCATTACCATCGTTTTCAAAGCTGCTCTCCAAA |
| AGGTTTTGAACGTCAAAAATGAAAGCGCTAATAAGCTGCTCTCCAAA |
| AATGGTTTACAACGCCAACATGTAGTTCAGCTAAGCTGCTCTCCAAA |
| AACCTACCGCGAATTATTCATTTCCAGTACATAAGCTGCTCTCCAAA |
| GCGTAAGAGAGAGCCAGCAGCAAAAAGGTTATAAGCTGCTCTCCAAA |
| GTTTGCCACCTCAGAGCCGCCACCGATACAGGAAGCTGCTCTCCAAA |
| GCCCAATACCGAGGAAACGCAATAGGTTTACCAAGCTGCTCTCCAAA |
| TAAGTCCTACCAAGTACCGCACTCTTAGTTGCAAGCTGCTCTCCAAA |
| CTGTAAATCATAGGTCTGAGAGACGATAAATAAAGCTGCTCTCCAAA |
| AGATTAGATTTAAAAGTTTGAGTACACGTAAAAAGCTGCTCTCCAAA |
| ATTGAGGGTAAAGGTGAATTATCAATCACCGGAAGCTGCTCTCCAAA |
| TCTTACCAGCCAGTTACAAAATAAATGAAATAAAGCTGCTCTCCAAA |
| AATTACTACAAATTCTTACCAGTAATCCCATCAAGCTGCTCTCCAAA |
| TTTAACGTTCGGGAGAAACAATAATTTTCCCTAAGCTGCTCTCCAAA |
| TAGCCCTACCAGCAGAAGATAAAAACATTTGAAAGCTGCTCTCCAAA |
| GCATAAAGTTCCACACAACATACGAAGCGCCAAAGCTGCTCTCCAAA |
| TTTCATTTGGTCAATAACCTGTTTATATCGCGAAGCTGCTCTCCAAA |
| CTCATCTTGAGGCAAAAGAATACAGTGAATTTAAGCTGCTCTCCAAA |
| GAATAGCCGCAAGCGGTCCACGCTCCTAATGAAAGCTGCTCTCCAAA |
| GGCGATCGCACTCCAGCCAGCTTTGCCATCAAAAGCTGCTCTCCAAA |
| ACCGTTCTAAATGCAATGCCTGAGAGGTGGCAAAGCTGCTCTCCAAA |
| GAAGCAAAAAAGCGGATTGCATCAGATAAAAAAAGCTGCTCTCCAAA |
| ACGAGTAGTGACAAGAACCGGATATACCAAGCAAGCTGCTCTCCAAA |
| ACTGCCCGCCGAGCTCGAATTCGTTATTACGCAAGCTGCTCTCCAAA |
| CTTTCATCCCCAAAAACAGGAAGACCGGAGAGAAGCTGCTCTCCAAA |
| CAATAAATACAGTTGATTCCCAATTTAGAGAGAAGCTGCTCTCCAAA |
| TTTGCCAGATCAGTTGAGATTTAGTGGTTTAAAAGCTGCTCTCCAAA |
| CGCCTGATGGAAGTTTCCATTAAACATAACCGAAGCTGCTCTCCAAA |
| TGGACTCCCTTTTCACCAGTGAGACCTGTCGTAAGCTGCTCTCCAAA |
| ATTAAGTTCGCATCGTAACCGTGCGAGTAACAAGCTGCTCTCCAAA |
| TCAGGTCACTTTTGCGGGAGAAGCAGAATTAGAAGCTGCTCTCCAAA |
| TTTTTGCGCAGAAAACGAGAATGAATGTTTAGAAGCTGCTCTCCAAA |
| CGATTTTAGAGGACAGATGAACGGCGCGACCTAAGCTGCTCTCCAAA |
| ACGCAAAGGTCACCAATGAAACCAATCAAGTTCATTAACATCATCAG |

| |
|---|
| GAGGCGTTAGAGAATAACATAAAAGAACACCCCATTAACATCATCAG |
| TTTTAGTTTTTCGAGCCAGTAATAAATTCTGTCATTAACATCATCAG |
| TGGATTATGAAGATGATGAAACAAAATTTCATCATTAACATCATCAG |
| GCCAACAGTCACCTTGCTGAACCTGTTGGCAACATTAACATCATCAG |
| TCGGCATTCCGCCGCCAGCATTGACGTTCCAGCATTAACATCATCAG |
| ATCAGAGAAAGAACTGGCATGATTTTATTTTGCATTAACATCATCAG |
| AATGCAGACCGTTTTTATTTTCATCTTGCGGGCATTAACATCATCAG |
| AAATCAATGGCTTAGGTTGGGTTACTAAATTTCATTAACATCATCAG |
| CTAAAATAGAACAAAGAAACCACCAGGGTTAGCATTAACATCATCAG |
| AGCGCCAACCATTTGGGAATTAGATTATTAGCCATTAACATCATCAG |
| TATTTTGCTCCCAATCCAAATAAGTGAGTTAACATTAACATCATCAG |
| AGGCGTTACAGTAGGGCTTAATTGACAATAGACATTAACATCATCAG |
| ACAGAAATCTTTGAATACCAAGTTCCTTGCTTCATTAACATCATCAG |
| GAATGGCTAGTATTAACACCGCCTCAACTAATCATTAACATCATCAG |
| AACCAGAGACCCTCAGAACCGCCAGGGGTCAGCATTAACATCATCAG |
| CTAATTTATCTTTCCTTATCATTCATCCTGAACATTAACATCATCAG |
| GGATTTAGCGTATTAAATCCTTTGTTTTCAGGCATTAACATCATCAG |
| CCGAAATCCGAAAATCCTGTTTGAAGCCGGAACATTAACATCATCAG |
| TTCGCCATTGCCGGAAACCAGGCATTAAATCACATTAACATCATCAG |
| AGACAGTCATTCAAAAGGGTGAGAAGCTATATCATTAACATCATCAG |
| TTTTAATTGCCCGAAAGACTTCAAAACACTATCATTAACATCATCAG |
| GAATAAGGACGTAACAAAGCTGCTCTAAAACACATTAACATCATCAG |
| GTGAGCTAGTTTCCTGTGTGAAATTTGGGAAGCATTAACATCATCAG |
| AAATAATTTTAAATTGTAAACGTTGATATTCACATTAACATCATCAG |
| TCAATTCTTTTAGTTTGACCATTACCAGACCGCATTAACATCATCAG |
| CCAAAATATAATGCAGATACATAAACACCAGACATTAACATCATCAG |
| GCGAAACATGCCACTACGAAGGCATGCGCCGACATTAACATCATCAG |
| AGTTTGGAGCCCTTCACCGCCTGGTTGCGCTCCATTAACATCATCAG |
| CAGCTGGCGGACGACGACAGTATCGTAGCCAGCATTAACATCATCAG |
| GGTAGCTAGGATAAAAATTTTTAGTTAACATCCATTAACATCATCAG |
| TACCTTTAAGGTCTTTACCCTGACAAAGAAGTCATTAACATCATCAG |
| TTTCAACTATAGGCTGGCTGACCTTGTATCATCATTAACATCATCAG |
| GCCAGCTGCCTGCAGGTCGACTCTGCAAGGCCATTAACATCATCAG |
| ACCCGTCGTCATATGTACCCCGGTAAAGGCTACATTAACATCATCAG |
| CAAAATTAAAGTACGGTGTCTGGAAGAGGTCACATTAACATCATCAG |
| ACTGGATAACGGAACAACATTATTACCTTATGCATTAACATCATCAG |
| GCTCCATGAGAGGCTTTGAGGACTAGGGAGTTCATTAACATCATCAG |
| CAAGCCCAATAGGAACCCATGTACAAACAGTT |
| AATGCCCCGTAACAGTGCCCGTATCTCCCTCA |
| TGCCTTGACTGCCTATTTCGGAACAGGGATAG |
| GAGCCGCCCCACCACCGGAACCGCGACGGAAA |
| TTATTCATAGGGAAGGTAAATATTCATTCAGT |
| CATAACCCGAGGCATAGTAAGAGCTTTTTAAG |

| |
|---|
| AAAAGTAATATCTTACCGAAGCCCTTCCAGAG |
| GCAATAGCGCAGATAGCCGAACAATTCAACCG |
| CCTAATTTACGCTAACGAGCGTCTAATCAATA |
| ATCGGCTGCGAGCATGTAGAAACCTATCATAT |
| GCGTTATAGAAAAAGCCTGTTTAGAAGGCCGG |
| GCTCATTTTCGCATTAAATTTTTGAGCTTAGA |
| TTAAGACGTTGAAAACATAGCGATAACAGTAC |
| TAGAATCCCTGAGAAGAGTCAATAGGAATCAT |
| CTTTTACACAGATGAATATACAGTAAACAATT |
| CGACAACTAAGTATTAGACTTTACAATACCGA |
| ACGAACCAAAACATCGCCATTAAATGGTGGTT |
| GAACGTGGCGAGAAAGGAAGGGAACAAACTAT |
| CGGCCTTGCTGGTAATATCCAGAACGAACTGA |
| CTCAGAGCCACCACCCTCATTTTCCTATTATT |
| CTGAAACAGGTAATAAGTTTTAACCCCTCAGA |
| AGTGTACTTGAAAGTATTAAGAGGCCGCCACC |
| GCCACCACTCTTTTCATAATCAAACCGTCACC |
| GACTTGAGAGACAAAAGGGCGACAAGTTACCA |
| GAAGGAAAATAAGAGCAAGAAACAACAGCCAT |
| ATTATTTAACCCAGCTACAATTTTCAAGAACG |
| GGTATTAAGAACAAGAAAAATAATTAAAGCCA |
| ACGCTCAAAATAAGAATAAACACCGTGAATTT |
| ATCAAAATCGTCGCTATTAATTAACGGATTCG |
| CCTGATTGAAAGAAATTGCGTAGACCCGAACG |
| TTATTAATGCCGTCAATAGATAATCAGAGGTG |
| AGGCGGTCATTAGTCTTTAATGCGCAATATTA |
| CCGCCAGCCATTGCAACAGGAAAAATATTTTT |
| CCCTCAGAACCGCCACCCTCAGAACTGAGACT |
| CCTCAAGAATACATGGCTTTTGATAGAACCAC |
| TAAGCGTCGAAGGATTAGGATTAGTACCGCCA |
| CACCAGAGTTCGGTCATAGCCCCCGCCAGCAA |
| AATCACCAAATAGAAAATTCATATATAACGGA |
| ATACCCAAGATAACCCACAAGAATAAACGATT |
| TTTTGTTTAAGCCTTAAATCAAGAATCGAGAA |
| CAAGCAAGACGCGCCTGTTTATCAAGAATCGC |
| CATATTTAGAAATACCGACCGTGTTACCTTTT |
| TAACCTCCATATGTGAGTGAATAAACAAAATC |
| GCGCAGAGATATCAAAATTATTTGACATTATC |
| ATTTTGCGTCTTTAGGAGCACTAAGCAACAGT |
| GCCACGCTATACGTGGCACAGACAACGCTCAT |
| GGAAATACCTACATTTTGACGCTCACCTGAAA |
| TATCACCGTACTCAGGAGGTTTAGCGGGGTTT |
| TGCTCAGTCAGTCTCTGAATTTACCAGGAGGT |

| |
|---|
| GGAAAGCGACCAGGCGGATAAGTGAATAGGTG |
| TGAGGCAGGCGTCAGACTGTAGCGTAGCAAGG |
| CCGGAAACACACCACGGAATAAGTAAGACTCC |
| TTATTACGGTCAGAGGGTAATTGAATAGCAGC |
| CTTTACAGTTAGCGAACCTCCCGACGTAGGAA |
| TCATTACCCGACAATAAACAACATATTTAGGC |
| AGAGGCATAATTTCATCTTCTGACTATAACTA |
| TATGTAAACCTTTTTTAATGGAAAAATTACCT |
| GAGCAAAAACTTCTGAATAATGGAAGAAGGAG |
| CGGAATTATTGAAAGGAATTGAGGTGAAAAAT |
| CTAAAGCAAGATAGAACCCTTCTGAATCGTCT |
| GAAATGGATTATTTACATTGGCAGACATTCTG |
| CCAGCAGGGGCAAAATCCCTTATAAAGCCGGC |
| GCTCACAATGTAAAGCCTGGGGTGGGTTTGCC |
| GCTTCTGGTCAGGCTGCGCAACTGTGTTATCC |
| GTTAAAATTTTAACCAATAGGAACCCGGCACC |
| AGGTAAAGAAATCACCATCAATATAATATTTT |
| TCGCAAATGGGGCGCGAGCTGAAATAATGTGT |
| AAGAGGAACGAGCTTCAAAGCGAAGATACATT |
| GGAATTACTCGTTTACCAGACGACAAAAGATT |
| CCAAATCACTTGCCCTGACGAGAACGCCAAAA |
| AAACGAAATGACCCCCAGCGATTATTCATTAC |
| CTTAAACATCAGCTTGCTTTCGAGCGTAACAC |
| TCGGTTTAGCTTGATACCGATAGTCCAACCTA |
| TGAGTTTCGTCACCAGTACAAACTTAATTGTA |
| CCCCGATTTAGAGCTTGACGGGGAAATCAAAA |
| GAGTTGCACGAGATAGGGTTGAGTAAGGGAGC |
| TCATAGCTACTCACATTAATTGCGCCCTGAGA |
| GAAGATCGGTGCGGGCCTCTTCGCAATCATGG |
| GCAAATATCGCGTCTGGCCTTCCTGGCCTCAG |
| TATATTTTAGCTGATAAATTAATGTTGTATAA |
| CGAGTAGAACTAATAGTAGTAGCAAACCCTCA |
| TCAGAAGCCTCCAACAGGTCAGGATCTGCGAA |
| CATTCAACGCGAGAGGCTTTTGCATATTATAG |
| AGTAATCTTAAATTGGGCTTGAGAGAATACCA |
| ATACGTAAAAGTACAACGGAGATTTCATCAAG |
| CAATGACACTCCAAAAGGAGCCTTACAACGCC |
| AAAAAAGGACAACCATCGCCCACGCGGGTAAA |
| TGTAGCATTCCACAGACAGCCCTCATCTCCAA |
| GTAAAGCACTAAATCGGAACCCTAGTTGTTCC |
| AGCTGATTACAAGAGTCCACTATTGAGGTGCC |
| CCCGGGTACTTTCCAGTCGGGAAACGGGCAAC |
| GTTTGAGGGAAAGGGGGATGTGCTAGAGGATC |

| |
|---|
| AGAAAAGCAACATTAAATGTGAGCATCTGCCA |
| CAACGCAATTTTTGAGAGATCTACTGATAATC |
| TCCATATACATACAGGCAAGGCAACTTTATTT |
| CAAAAATCATTGCTCCTTTTGATAAGTTTCAT |
| AAAGATTCAGGGGGTAATAGTAAACCATAAAT |
| CCAGGCGCTTAATCATTGTGAATTACAGGTAG |
| TTTCATGAAAATTGTGTCGAAATCTGTACAGA |
| ATATATTCTTTTTCACGTTGAAAATAGTTAG |
| AATAATAAGGTCGCTGAGGCTTGCAAAGACTT |
| CGTAACGATCTAAAGTTTTGTCGTGAATTGCG |
| ACCCAAATCAAGTTTTTTGGGGTCAAAGAACG |
| TGGTTTTTAACGTCAAAGGGCGAAGAACCATC |
| CTTGCATGCATTAATGAATCGGCCCGCCAGGG |
| TAGATGGGGGGTAACGCCAGGGTTGTGCCAAG |
| CATGTCAAGATTCTCCGTGGGAACCGTTGGTG |
| CTGTAATATTGCCTGAGAGTCTGGAAAACTAG |
| TGCAACTAAGCAATAAAGCCTCAGTTATGACC |
| AAACAGTTGATGGCTTAGAGCTTATTTAAATA |
| ACGAACTAGCGTCCAATACTGCGGAATGCTTT |
| CTTTGAAAAGAACTGGCTCATTATTTAATAAA |
| ACGGCTACTTACTTAGCCGGAACGCTGACCAA |
| AAAGGCCGAAAGGAACAACTAAAGCTTTCCAG |
| GAGAATAGCTTTTGCGGGATCGTCGGGTAGCA |
| ACGTTAGTAAATGAATTTTCTGTAAGCGGAGT |
| AACATCACTTGCCTGAGTAGAAGAACT |
| TGTAGCAATACTTCTTTGATTAGTAAT |
| AGTCTGTCCATCACGCAAATTAACCGT |
| ATAATCAGTGAGGCCACCGAGTAAAAG |
| ACGCCAGAATCCTGAGAAGTGTTTTT |
| TTAAAGGGATTTTAGACAGGAACGGT |
| AGAGCGGGAGCTAAACAGGAGGCCGA |
| TATAACGTGCTTTCCTCGTTAGAATC |
| GTACTATGGTTGCTTTGACGAGCACG |
| GCGCTTAATGCGCCGCTACAGGGCGC |
